# Biofilm hydrophobicity in environmental isolates of *Bacillus subtilis*

**DOI:** 10.1101/2021.04.29.441976

**Authors:** Margarita Kalamara, James C. Abbott, Cait E. MacPhee, Nicola. R. Stanley-Wall

## Abstract

Biofilms are communities of bacteria that are attached to a surface and surrounded by an extracellular matrix. The extracellular matrix protects the community from stressors in the environment, making biofilms robust. The Gram-positive soil bacterium *Bacillus subtilis*, particularly the isolate NCIB 3610, is widely used as a model for studying biofilm formation. *B. subtilis* NCIB 3610 forms colony biofilms that are architecturally complex and highly hydrophobic. The hydrophobicity is linked, in part, to the localisation of the protein BslA at the surface of the biofilm, which provides the community with increased resistance to biocides. As most of our knowledge about *B. subtilis* biofilm formation comes from one isolate, it is unclear if biofilm hydrophobicity is a widely distributed feature of the species. To address this knowledge gap, we collated a library of *B. subtilis* soil isolates and acquired their whole genome sequences. We used our new isolates to examine biofilm hydrophobicity and found that, although BslA is encoded and produced by all isolates in our collection, hydrophobicity is not a universal feature of *B. subtilis* colony biofilms. To test whether the matrix exopolymer poly γ-glutamic acid could be masking hydrophobicity in our hydrophilic isolates, we constructed deletion mutants and found, contrary to our hypothesis, that the presence of poly γ-glutamic acid was not the reason behind the observed hydrophilicity. This study highlights the natural variation in the properties of biofilms formed by different isolates and the importance of using a more diverse range of isolates as representatives of a species.

**Repositories:** Raw sequence reads and annotated assemblies have been submitted to the European Nucleotide Archive under accession PRJEB43128.

## Introduction

Biofilms are social communities of bacteria that are enveloped within a self-produced extracellular matrix. The biofilm matrix consists of exopolymers of various forms, secreted proteins, and extracellular DNA (1). This complex biomaterial provides the community with structure and protection from environmental threats. Threats that biofilms show increased resistance to include ultraviolet radiation, host immune responses, antibiotics, biocides, heat, oxidation, metal toxicity, and physical forces (2). Thus, biofilm formation can be used as a survival mechanism by bacteria, allowing them to colonise diverse niches and persist in hostile environments.

*Bacillus subtilis* is a Gram-positive bacterium. The undomesticated isolate NCIB 3610 is the progenitor of the laboratory strain 168 (3), and has been used extensively for researching the regulatory mechanisms of biofilm formation and to uncover the materials used in the matrix (4, 5). The main biofilm matrix components of NCIB 3610 are the protein fibres formed by TasA and TapA (6), an exopolysaccharide (EPS) synthesised by the products of the *epsA-O* operon (7), and the secreted protein BslA (8). While the study of biofilm matrix composition in different isolates of the species has so far been limited, a study on six environmental isolates of *B. subtilis* reported that the matrix components TasA and EPS are conserved and play a crucial role in biofilm formation (9). Further studies have demonstrated a varied reliance on the exopolymer poly γ-glutamic acid (γ-PGA) in environmental *B. subtilis* isolates (10, 11). γ-PGA is correlated with a mucoid colony phenotype (12), and is a major component of the biofilm matrix in selected isolates, contributing to complex biofilm colony and pellicle architecture as well as plant root attachment (10, 11). In the reference isolate NCIB 3610, deletion of the genomic region involved in γ-PGA biosynthesis has an impact on biofilm architecture only under specific environmental conditions (10, 13, 14). Taken together, these findings suggest a difference in the presence, distribution and / or production of matrix exopolymeric substances in different isolates of the species.

A remarkable property of colony and pellicle biofilms formed by the model isolate NCIB 3610 is the production of a highly water repellent coating (15). This property provides an important protection mechanism for the resident bacteria, imparting increased resistance to biocides, gas penetration and solvents (15, 16). The dominant protein responsible for hydrophobicity is BslA, which works synergistically with the EPS of the matrix to form an elastic layer around the multicellular community. As such, mutations in the *bslA* gene result in a biofilm-deficient strain (17–19) with a hydrophilic phenotype. BslA has an immunoglobulin-like fold in which the loops at one end form a “cap” region made up of hydrophobic residues. The protein can be found in two conformations: “cap in” or “cap out”. In an aqueous environment, the hydrophobic residues are hidden in the interior of the protein (“cap in”), while when at the surface or an interface, the hydrophobic residues are exposed, resulting in the “cap out” conformation (18, 19). This conformational flexibility allows BslA to persist in the aqueous environment of the biofilm matrix, but also confer hydrophobicity when at the biofilm–air interface (18, 19). Further studies into the mechanisms by which BslA provides the biofilm with surface hydrophobicity revealed the importance of two cysteine residues at the C-terminus of the protein (C178 and C180), termed the “CxC” motif (16). The cysteines of the CxC motif form intermolecular disulphide bonds resulting in dimerization of the BslA monomers. Interestingly, although the BslA dimerization is crucial for biofilm hydrophobicity, monomers of the protein are sufficient to give rise to a complex colony morphology indistinguishable from the wild type biofilm (16). Another variable that contributes to biofilm hydrophobicity is biofilm structure. Growth of the model isolate NCIB 3610 under different conditions resulted in variations in colony structure and level of hydrophobicity (20). Finally, a study has additionally reported that the presence of metal ions Cu and Zn can render the NCIB 3610 biofilms hydrophilic. It was demonstrated that hydrophilicity due to the presence of these ions increased the biofilm’s susceptibility to antibiotic treatment, further strengthening the evidence for the protective nature of biofilm hydrophobicity (21).

While the molecular mechanism by which BslA functions to provide the community with hydrophobicity and consequently protection from environmental threats is understood, it is unknown how broadly this property is conserved among different isolates of species. To address this knowledge gap, here we used a citizen science approach to assemble a library of 39 environmental isolates of *B. subtilis* that were extracted from soil. We sequenced the isolates and examined the presence and conservation of BslA. All isolates were found to encode BslA and their DNA and protein sequences show strong sequence conservation. We tested colony biofilm hydrophobicity and found that only a minority of the isolates in our collection were hydrophobic under the conditions tested, despite BslA being produced in the mature biofilms of both hydrophobic and hydrophilic isolates alike. To test whether the mucoid polymer γ-PGA masked hydrophobicity in the hydrophilic isolates, deletion mutants were constructed in a selected subset of isolates. Our results show that presence of γ-PGA is not responsible for biofilm hydrophilicity. Thus, the reason why hydrophobicity is not conserved, despite BslA being encoded and produced by all isolates, remains unclear. Taken together, our findings illustrate the importance of BslA for hydrophobicity in colony biofilms but uncover further diversity in the biofilm matrices produced across the species.

## Materials and Methods

### Bacterial strains and growth conditions

All *B. subtilis* isolates used in this study are listed in Table 1. The strains were routinely grown on lysogeny broth (LB: 1% (w/v) Bacto-peptone, 1% (w/v) NaCl, 0.5% (w/v) yeast extract and 1.5% (w/v) agar) plates or in liquid cultures at 37°C. For biofilm experiments, the strains were grown on MSgg agar plates (5 mM potassium phosphate (pH 7), 100 mM MOPS (pH=7), 2 mM MgCl_2_, 700 μM CaCl_2_, 50 μM MnCl_2_, 50 μM FeCl_3_, 1 μM ZnCl_2_, 2 μM thiamine 0.5% (v/v) glycerol, 0.5% (w/v) glutamate, 1.5% (w/v) agar) (7). For competency assays, an altered version of 10 x Modified Competency (MC) media was used (10.7 g K_2_HPO_4_, 5.2 g KH_2_PO_4_, 20 g dextrose, 0.88 g sodium citrate dehydrate, 2.2 g L-glutamic acid monopotassium salt, and 1 g tryptone per 100 ml) (22). Antibiotics were added as required at the following concentrations: spectinomycin (100 μg/ml); chloramphenicol (5 μg/ml); kanamycin (10 μg/ml).

**Table 1:** Strains used in this study.

| Strain | Species | Genotype <sup>a</sup> | Source <sup>b</sup> |
| --- | --- | --- | --- |
| NCIB 3610 | *B. subtilis* | Wild type | B.G.S.C. |
| NRS2097 | *B. subtilis* | NCIB 3610 *bslA::cat*<sup>a</sup> | (32) |
| NRS6084 | *B. subtilis* | Wild type (Blairgowrie, UK) | This study |
| NRS6085 | *B. subtilis* | Wild type (Cromarty Firth, UK, leaf litter) | This study |
| NRS6092 | *B. subtilis* | Wild type (Tayport, UK, homemade compost) | This study |
| NRS6094 | *B. subtilis* | Wild type (Tayport, UK, garden soil) | This study |
| NRS6096 | *B. subtilis* | Wild type (Tayport, UK, garden soil) | This study |
| NRS6099 | *B. subtilis* | Wild type (Tayport, UK, garden soil) | This study |
| NRS6103 | *B. subtilis* | Wild type (Tayport, UK, community garden soil) | This study |
| NRS6105 | *B. subtilis* | Wild type (Tayport, UK, community garden soil) | This study |
| NRS6107 | *B. subtilis* | Wild type (Tayport, UK, garden soil) | This study |
| NRS6108 | *B. subtilis* | Wild type (Tayport, UK, garden soil) | This study |
| NRS6110 | *B. subtilis* | Wild type (Tayport, UK, vegetable plot) | This study |
| NRS6111 | *B. subtilis* | Wild type (Tayport, UK, vegetable plot) | This study |
| NRS6116 | *B. subtilis* | Wild type (Tayport, UK, garden soil) | This study |
| NRS6118 | *B. subtilis* | Wild type (Tayport, UK, potato patch) | This study |
| NRS6120 | *B. subtilis* | Wild type (Tayport, UK, potato patch) | This study |
| NRS6121 | *B. subtilis* | Wild type (Tayport, UK, shrub bed) | This study |
| NRS6127 | *B. subtilis* | Wild type (Tayport, UK, garden soil) | This study |
| NRS6128 | *B. subtilis* | Wild type (Tayport, UK, garden soil) | This study |
| NRS6131 | *B. subtilis* | Wild type (Tayport, UK, worm bin) | This study |
| NRS6132 | *B. subtilis* | Wild type (Tayport, UK, worm bin) | This study |
| NRS6134 | *B. subtilis* | Wild type (Tayport, UK, vegetable patch) | This study |
| NRS6137 | *B. subtilis* | Wild type (Tayport, UK, community garden soil) | This study |
| NRS6141 | *B. subtilis* | Wild type (Tayport, UK, garden soil) | This study |
| NRS6145 | *B. subtilis* | Wild type (Tayport, UK, garden soil) | This study |
| NRS6148 | *B. subtilis* | Wild type (Tayport, UK, garden soil) | This study |
| NRS6153 | *B. subtilis* | Wild type (Tayport, UK, garden soil) | This study |
| NRS6160 | *B. subtilis* | Wild type (Tayport, UK, garden soil) | This study |
| NRS6167 | *B. subtilis* | Wild type (Tayport, UK, soil from planter) | This study |
| NRS6181 | *B. subtilis* | Wild type (Lochee, UK, garden soil) | This study |
| NRS6183 | *B. subtilis* | Wild type (Lochee, UK, garden soil) | This study |
| NRS6185 | *B. subtilis* | Wild type (Lochee, UK, garden soil) | This study |
| NRS6186 | *B. subtilis* | Wild type (Newport, UK, garden soil) | This study |
| NRS6187 | *B. subtilis* | Wild type (Newport, UK, garden soil) | This study |
| NRS6190 | *B. subtilis* | Wild type (Tayport, UK, garden soil) | This study |
| NRS6194 | *B. subtilis* | Wild type (Tayport, UK, garden soil) | This study |
| NRS6202 | *B. subtilis* | Wild type (Kirriemuir, UK, garden soil) | This study |
| NRS6204 | *B. subtilis* | Wild type (Kirriemuir, UK, garden soil) | This study |
| NRS6205 | *B. subtilis* | Wild type (Kirriemuir, UK, garden soil) | This study |
| NRS6206 | *B. subtilis* | Wild type (Kirriemuir, UK, garden soil) | This study |
| NRS6901 | *B. subtilis* | NRS6105 *pgsB::spec* | pNW2301 into NRS6105 |
| NRS6902 | *B. subtilis* | NRS6153 *pgsB::spec* | pNW2301 into NRS6153 |
| NRS6903 | *B. subtilis* | NRS6096 *pgsB::spec* | pNW2301 into NRS6096 |
| NRS6904 | *B. subtilis* | NRS6118 *pgsB::spec* | pNW2301 into NRS6118 |
| NRS7203 | *B. subtilis* | NRS6105 *bslA::kan* | pNW2305 into NRS6105 |
| NRS7204 | *B. subtilis* | NRS6153 *bslA::kan* | pNW2305 into NRS6153 |
| NRS7205 | *B. subtilis* | NRS6127 *bslA::kan* | pNW2305 into NRS6127 |
| NRS7206 | *B. subtilis* | NRS6145 *bslA::kan* | pNW2305 into NRS6145 |

**a** The abbreviation “spec” indicates spectinomycin resistance; “cat” indicates chloramphenicol resistance and “kan” kanamycin resistance.
**b** The method of strain construction is indicated with the plasmid inserted into the parental strain.

### Isolating bacteria through citizen science

A citizen science approach was used for isolating bacteria from soil. Participants brought soil from their gardens to the citizen science events. 1 g of soil was mixed with 10 ml of sterile water and the soil and water mixture was serially diluted. Approximately 100 μl of the 10^-1^ and 10^-2^ dilutions were plated on LB plates supplemented with 100 μg/ml chlorhexidine to inhibit fungal growth (these were labelled “diversity” plates). The combined water and soil solution was incubated in an 80 °C water-bath for 10 min to kill vegetative cells, enriching endospore forming bacteria. 100 μl of the heat-treated samples were subsequently plated onto LB plates supplemented with 100 μg/ml chlorhexidine plates to isolate spore forming bacteria. The diversity plates were incubated at room temperature for approximately one week before imaging. The heat-treated sample plates were incubated at 30 °C overnight. The next day the plates were imaged, and three colonies were chosen from each plate. The selected colonies were streak purified twice before storing as glycerol stocks at −80 °C.

### Species classification

Colony PCR and sequencing of a partial 16S rRNA region covering V3-V5 (740 bp) were used for preliminary species classification. For DNA extraction, single colonies of the stocked isolates were re-suspended in 50 μl of sterile water and incubated at −80 °C for 10 min. The samples were immediately moved to 95 °C and incubated for a further 5 min. The samples were centrifuged and 5 μl of the supernatant was used as a template for the PCR. The PCR reactions were performed in a final volume of 25 μl, using of 12.5 μl of GoTaq^®^ Green Master Mix (1x). The primers used were 338f (stocked as NSW2750) 5’-TCACGRCACGAGCTGACGAC-3’ and 1061r (stocked as NRW2751) 5’-ACTCCTACGGGAGGCAGC-3’ and were added to the reaction at a final concentration of 0.4 μM each. After PCR amplification, the fragments were sent for sequencing using the same primers as those used for colony PCR. Sequence identity assessment was performed on the sequences retrieved using BLASTn (23). Isolates that were preliminarily classified as *B. subtilis* were sent for whole genome sequencing.

### Whole genome sequencing

Genome sequencing was provided by MicrobesNG (http://www.microbesng.uk). For sample preparation, single colonies of each strain to be sequenced were re-suspended in sterile PBS buffer and streaked onto LB agar plates. The plates were incubated at 37 °C overnight and the following day, the cells were harvested, placed into the barcoded bead tubes provided and sent to the MicrobesNG facilities. There, for each sample, three beads were washed with extraction buffer containing lysozyme and RNase A, incubated for 25 min at 37 °C. Proteinase K and RNaseA were added and incubated for 5 min at 65 °C. Genomic DNA was purified using an equal volume of SPRI beads and resuspended in EB buffer. DNA was quantified in triplicate with the Quantit dsDNA HS assay in an Eppendorf AF2200 plate reader. Genomic DNA libraries were prepared using Nextera XT Library Prep Kit (Illumina, San Diego, USA) following the manufacturer’s protocol with the following modifications: two nanograms of DNA instead of one were used as input, and PCR elongation time was increased to 1 min from 30 s. DNA quantification and library preparation were carried out on a Hamilton Microlab STAR automated liquid handling system. Pooled libraries were quantified using the Kapa Biosystems Library Quantification Kit for Illumina on a Roche light cycler 96 qPCR machine. Libraries were sequenced on the Illumina HiSeq using a 250 bp paired end protocol. Reads were adapter trimmed using Trimmomatic 0.30 with a sliding window quality cutoff of Q15 (24). De novo assembly was performed on samples using SPAdes version 3.7 (25), and contigs were annotated using Prokka 1.11 (26). Annotated draft assemblies of the sequencing results were acquired and whole genome sequencing data were visualised in Artemis software (27). Raw sequence reads and annotated assemblies have been submitted to the European Nucleotide Archive under accession PRJEB43128.

### Phylogenetic tree construction

The nucleotide sequences of *gyrA, rpoB, dnaJ* and *recA* were extracted and concatenated. The same sequences for the reference strains were retrieved from NCBI, concatenated, and included in the analysis (Table S1). The sequences were aligned in Jalview (28) by MAFFT using the G-INS-I algorithm and MEGA7 software (29) was used to construct a maximum likelihood phylogenetic tree with 100 bootstrap repeats. The resulting tree was rooted on *B. amyloliquefaciens*, which was included in the analysis as an outgroup.

### BslA alignment

The whole genome sequences of soil isolates were visualized in Artemis software and the nucleotide and amino acid sequences of BslA were extracted. The same sequences for the model NCIB3610 were retrieved from NCBI. Jalview (28) was used to align, annotate, wrap, and save the sequence alignment as a TIFF file.

### Screening for genetic competency

Genetic competency assays were performed as described by Konkol *et al*., 2013 (22). Briefly, a 2 ml culture of each isolate to be transformed was set up in 1x MC media supplemented with 3 mM of MgSO_4_ and 875 μM of FeCl_3_ and grown for 4.5 h at 37 °C with gentle agitation. A 400 μl aliquot of each culture was then mixed with 25 μl of plasmid pBL165 (30). This plasmid carries *gfpmut2* (encoding a variant of GFP) linked to a chloramphenicol resistance cassette (*cat*) and flanked by the 5’ and 3’ coding regions of *amyE*, allowing for integration of the *gfpmut2* and *cat* construct into the *amyE* locus upon successful transformation. After addition of the plasmid, the cultures were incubated 37 °C for an additional 90 min before plating onto selective media (LB containing 5 μg/ml chloramphenicol). The plates were incubated at 37 °C overnight and transformants were screened for GFP production using fluorescence imaging and disruption of amylase activity using a potato starch assay (31).

### Strain construction

For construction of the *pgsB* and *bslA* mutants, plasmids (pNW2301 and pNW2305 respectively) were synthetically produced by Genscript ™. The sequences used are shown in Table S2 and the background vector used was pUC57. The acquired plasmids were transformed into the selected isolates using the same method as described for the genetic competency assays above, changing the antibiotic used for selection to spectinomycin or kanamycin as required. Disruption of the *pgsB* gene was assessed using primers NSW2763 (5’-GTTAGAGAATTCGGACTCGTATG-3’) and NSW2765 (5’-CAAGAAATGGTACCGTGGAATC-3’) which bind 500 bp upstream of the *pgsB* start codon and 600 bp from the start of the SpecR cassette, respectively. Disruption of the *bslA* gene was assessed using primers NSW2776 (5’-GTATGGATCCGACGCTTGACGAAATGC-3’) and NSW2769 (5’-GCACTCCGCATACAGCTCG-3’) which bind 600 bp upstream of *bslA* start codon and 520 bp from the start of the KanR cassette, respectively.

### Biofilm morphology assays

*B. subtilis* isolates were streaked out on LB agar plates and incubated at 30 °C overnight. The following day, single colonies were grown in 3 ml of LB broth at 37 °C with agitation. The cultures were grown to an OD_600_ of 1 and 10 μl of the cultures were spotted onto MSgg media plates. The plates were incubated at 30 °C for 48 h before imaging. Biofilm imaging was performed using a Leica MZ16 FA stereoscope and LAS version 2.7.1.

### Biofilm hydrophobicity assays

Biofilm hydrophobicity was tested by measuring the contact angle between the surfaces of biofilms grown at 30 °C for 48 h and a 10 μl drop of water, as described previously (19). The measurements were taken 5 min after initial placement of the water droplet on the biofilm surface using a ThetaLite TL100 optical tensiometer. Contact angles were determined with OneAttension, using the Young-Laplace equation. Contact angles above 90° are indicative of a hydrophobic surface, whereas surfaces with contact angles below 90° are considered hydrophilic. A minimum of three biological and three technical replicates were performed for each isolate.

### Biofilm protein extraction

48-hour old biofilms were removed from agar plates using sterile loops and placed in 250 μl of BugBuster (Novagen). The biofilm was disrupted by repetitive passage through a sterile 23-gauge needle. The samples were gently sonicated and incubated at 26 °C for 20 min with gentle agitation. The samples were centrifuged at 17,000 × g for 10 min and the supernatant was kept and analysed by immunoblot after separation by SDS-PAGE.

### Immunoblot analysis

Biofilm protein extracts were separated using 14% (w/v) SDS-PAGE. The proteins were transferred onto a PVDF membrane by electroblotting at 100 mA for 75 min. The membrane was incubated in TBS (20 mM TrisoHCl (pH 8.0), 0.15 M NaCl) supplemented with 3% (w/v) skimmed milk powder at 4 °C overnight with agitation. The next day the membrane was washed with TBS-T (TBS containing 0.05% (v/v) Tween 20) and incubated in TBS-T containing 3% (w/v) skimmed milk powder with purified anti-BslA antibody (32) at a 1:500 (v/v) dilution for 2 h at room temperature with shaking. After washing with TBS-T, the membrane was incubated for 45 min in TBS-T with 3% (w/v) skimmed milk powder with a goat anti-rabbit secondary antibody, conjugated to horseradish peroxidase, at a 1:5,000 dilution at room temperature. The membrane was washed with TBS-T, developed by the addition of ECL peroxidase reagent and visualised using an X-ray film.

## Results

### Collating a library of *Bacillus subtilis* soil isolates through citizen science

To test biofilm hydrophobicity in a range of natural *B. subtilis* isolates, we collated a library of environmental *B. subtilis* strains. We took a citizen science approach to acquire the isolates, engaging members of our local community with microbiology research, while simultaneously obtaining the specimens. To do this, outreach events were hosted in conjunction with a local community garden, and participants were guided through the actions of processing and plating soil samples for *B. subtilis* isolation. We prepared “diversity” plates, to show the range of bacteria that can be isolated from soil, and “*Bacillus*” plates, to select for endospore forming bacteria after heat treatment of the soil samples (Figure 1A). These steps were performed in the field by the participants. Following incubation, the plates were imaged and colonies of 135 endospore-forming bacteria were isolated and purified in the laboratory. All the purified isolates were preliminarily taxonomically classified based on 16S rRNA sequencing (Table S3). 41 of the 135 stocked isolates were classified as *B. subtilis* using this approach. Other species preliminarily classified included other commonly isolated soil bacteria such as *B. amyloliquefaciens, Lysinibacillus fusiformis, Lysinibacillus parviboronicapiens* as examples. Each participant received images of the plates they had prepared and a report outlining the different bacterial species found in their soil samples, as well as some information of their roles in the soil ecosystem.

**Figure 1:**
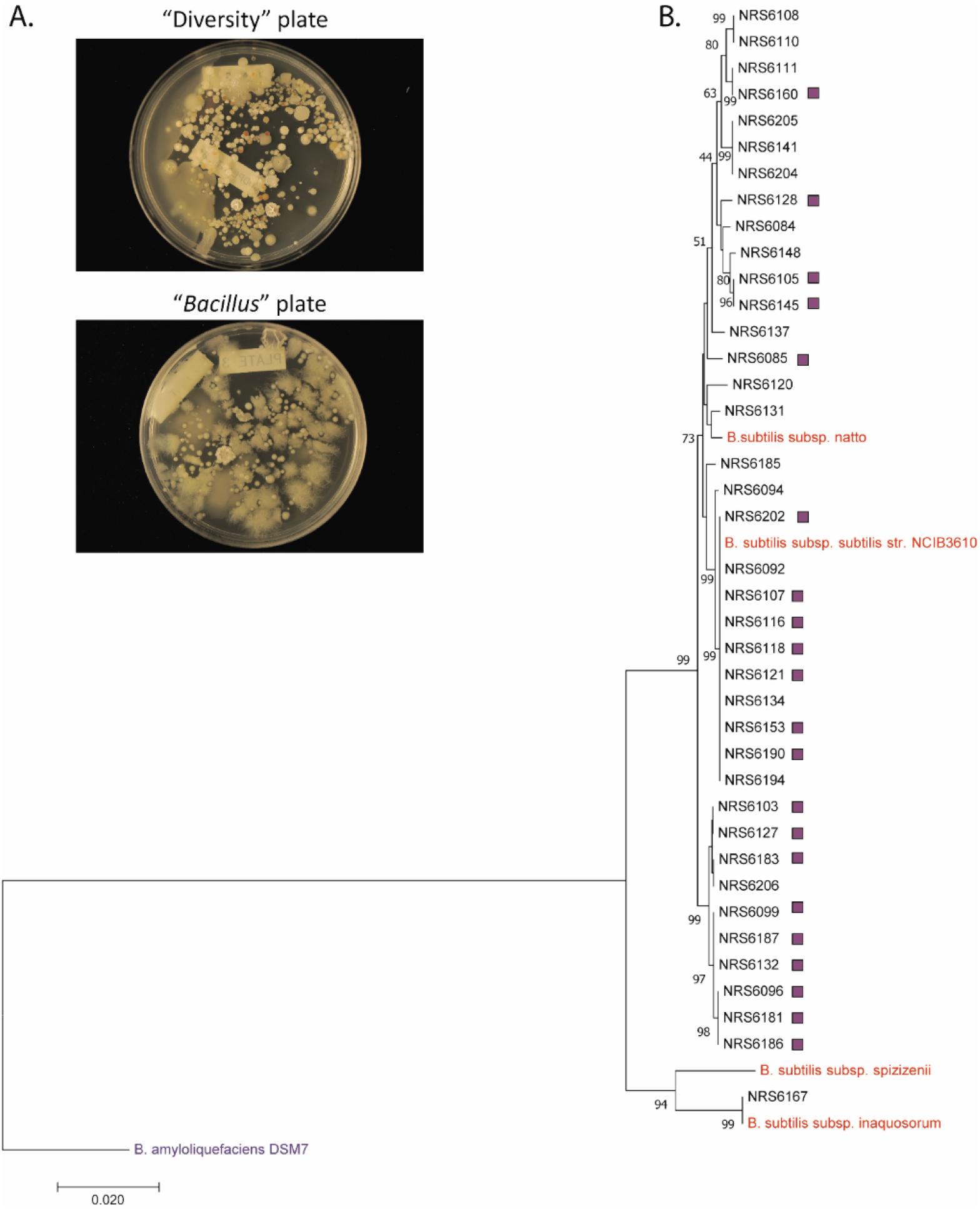
Soil isolates of *B. subtilis*. (**A**) Images of two example plates produced during the citizen science workshops. “Diversity” plate refers to the soil and water samples plated before heat treatment and the “*Bacillus*” plate is the result of plating after heat treatment to select for endospore forming bacteria. (**B**) Maximum likelihood phylogenetic tree based on the sequences of *gyrA_rpoB_dnaJ_recA*. “NRS” isolates are those acquired in this study. Sequences for reference *B. subtilis* strains (in red) and closely related *Bacillus* strain, *B. amyloliquefaciens* (in blue), were retrieved from NCBI (See Table S1). Genetically competent isolates are indicated with the purple square to the right of the strain name.

### Phylogenetic analysis of *B. subtilis* soil isolates

The 41 isolates preliminarily classified as *B. subtilis* were sent for whole genome sequencing. 39 of the 41 strains were confirmed to belong to the *B. subtilis* species, while the remaining 2 were identified as closely related species in the *B. subtilis* clade, namely *B. amyloliquefaciens* and *B. methylotrophicus* (see Table S3). The average genome size of the 39 *B. subtilis* isolates was approximately 4.2 Mbp with a range of 3.97 Mbp to 4.32 Mbp, and comprised an average GC content of 43.49%, with a minimum of 43.13% and a maximum of 43.9% recorded (Table S4). To explore the relatedness between the novel isolates a phylogenetic tree was constructed. Reference isolates belonging to different *B. subtilis* subspecies (*inaquosorum, subtilis* and *spizizenii*) were included in the analysis (Table S1) to allow for a more detailed assessment of the evolutionary relationships amongst isolates. A maximum-likelihood tree was constructed based on the concatenated sequences of four housekeeping genes (*gyrA, rpoB, dnaJ, recA*) with 100 bootstrap repeats (Figure 1B). Most of the environmental isolates were more closely related to *B. subtilis subsp. subtilis*, except for isolate NRS6167, which clustered with *B. subtilis subsp. inaquosorum*.

### BslA is present and conserved in all isolates of *B. subtilis*

The secreted protein BslA is linked to the non-wetting biofilm phenotype of *B. subtilis* NCIB 3610 colony and pellicle biofilms (8). To start to explore the generality of colony biofilm hydrophobicity of our new isolates, we examined the presence and conservation of *bslA* at a genomic level. We compared both the *bslA* nucleotide and the BslA amino acid sequences of the soil isolates and the model isolate NCIB 3610 (28). BslA was encoded by each of the isolates and was well conserved, with 38 out of 39 isolates having a sequence that was 100% identical to that of NCIB 3610 at the amino acid level. The only isolate that showed variation in the BslA sequence was the most distantly related to the rest based on phylogenetic analysis (NRS6167) (Figure 2A). The differences between the amino acid sequence of BslA from NRS6167 and the rest of the isolates were not present in regions of BslA known to be needed for function in NCIB 3610 (namely the cap regions and the CxC motif) (16, 19). Consistent with the conservation of the amino acid sequence, the *bslA* nucleotide sequences showed limited variability, with all isolates sharing a *bslA* nucleotide identity of 94.3-100% to that of NCIB 3610 (Figure S1).

**Figure 2:**
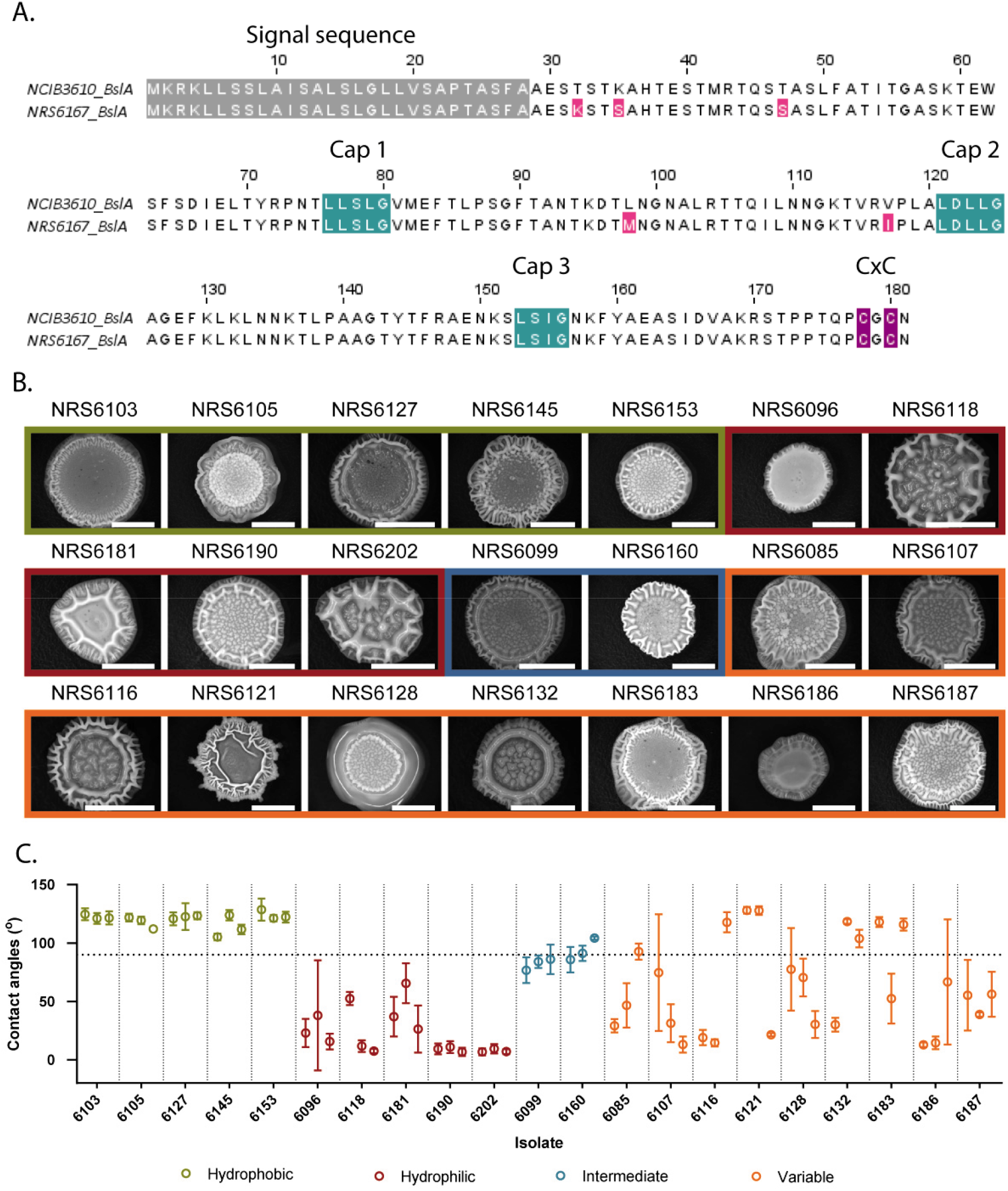
Biofilm morphology and hydrophobicity in environmental isolates of *B. subtilis*. (**A**) Alignment of BslA encoded by NCIB 3610 and that of the soil isolate NRS6167. The predicted signal sequences are highlighted in grey. The highlighted regions represent the conserved cap regions in the monomeric structure (19) (turquoise) and “CxC” motif (16) (purple), previously shown to be required for biofilm hydrophobicity. Amino acid residues highlighted in pink represent variations from the NCIB 3610 sequence identified in the environmental strain. (**B**) Representative images of 48 h biofilms formed by the soil isolates in this study. The scale bars represent 1 cm. (**C**) Biofilm hydrophobicity assay results of *B. subtilis* isolates. Results represent the mean value for each of three biological repeats (shown as the three points per isolate on the graph). Error bars represent the standard deviation of three technical repeats. The horizontal line represents the 90° contact angle cut-off for hydrophobicity and the vertical lines separate the data for each of the isolates. The four different colours of the data points represent the different phenotypes, as described in the legend below the graph. The coloured borders in (**B**) correspond to the colour coded hydrophobicity phenotypes in (**C**). The values of the parental strains are the same as show in Fig. 3B and 4B and are repeated for clarity (Table S5).

### Biofilm hydrophobicity is not a conserved feature of the *B. subtilis* biofilm

As BslA facilitates hydrophobicity in NCIB 3610, the strong conservation of the BslA sequences led us to hypothesize that all isolates would form colony biofilms with non-wetting hydrophobic upper surfaces. We reasoned that strains with natural genetic competence would be beneficial for further studies and would allow, for example, the generation of deletion strains. We therefore screened all isolates for natural genetic competency using an integrative plasmid with a selective marker. We eliminated strains that were not naturally genetically tractable and one further isolate (NRS6167) that was found to be resistant to the antibiotic used for selection of successful transformants. 21 out of the remaining 38 isolates in our library were genetically competent and used in further experimental work (Figure 1B).

Colony biofilm hydrophobicity assays were conducted on the 21 genetically competent soil isolates using NCIB 3610 as a reference. The contact angle between the surface of the biofilm and a drop of water was calculated to determine wetting and non-wetting surfaces. All isolates formed structured biofilms under laboratory conditions and displayed a variety of different morphologies (Figure 2B). Hydrophobicity was only consistently observed in five of the isolates tested, while another five of the isolates in our collection were consistently hydrophilic. Two of the remaining isolates had a borderline hydrophobic phenotype and the others were highly variable in terms of the contact angle measured (Figure 2C and Table S5). Therefore, we conclude that among the 21 isolates of *B. subtilis* in our collection, biofilm hydrophobicity is not a conserved feature. For four of the consistently hydrophobic isolates, BslA was linked as a causative agent of the hydrophobicity (and biofilm architecture) since *bslA* deletion strains resulted in an altered biofilm phenotype (Figure 3A) and a hydrophilic surface (Figure 3B). We were unable to obtain a *bslA* deletion strain for the remaining hydrophobic isolate (strain NRS6103).

**Figure 3:**
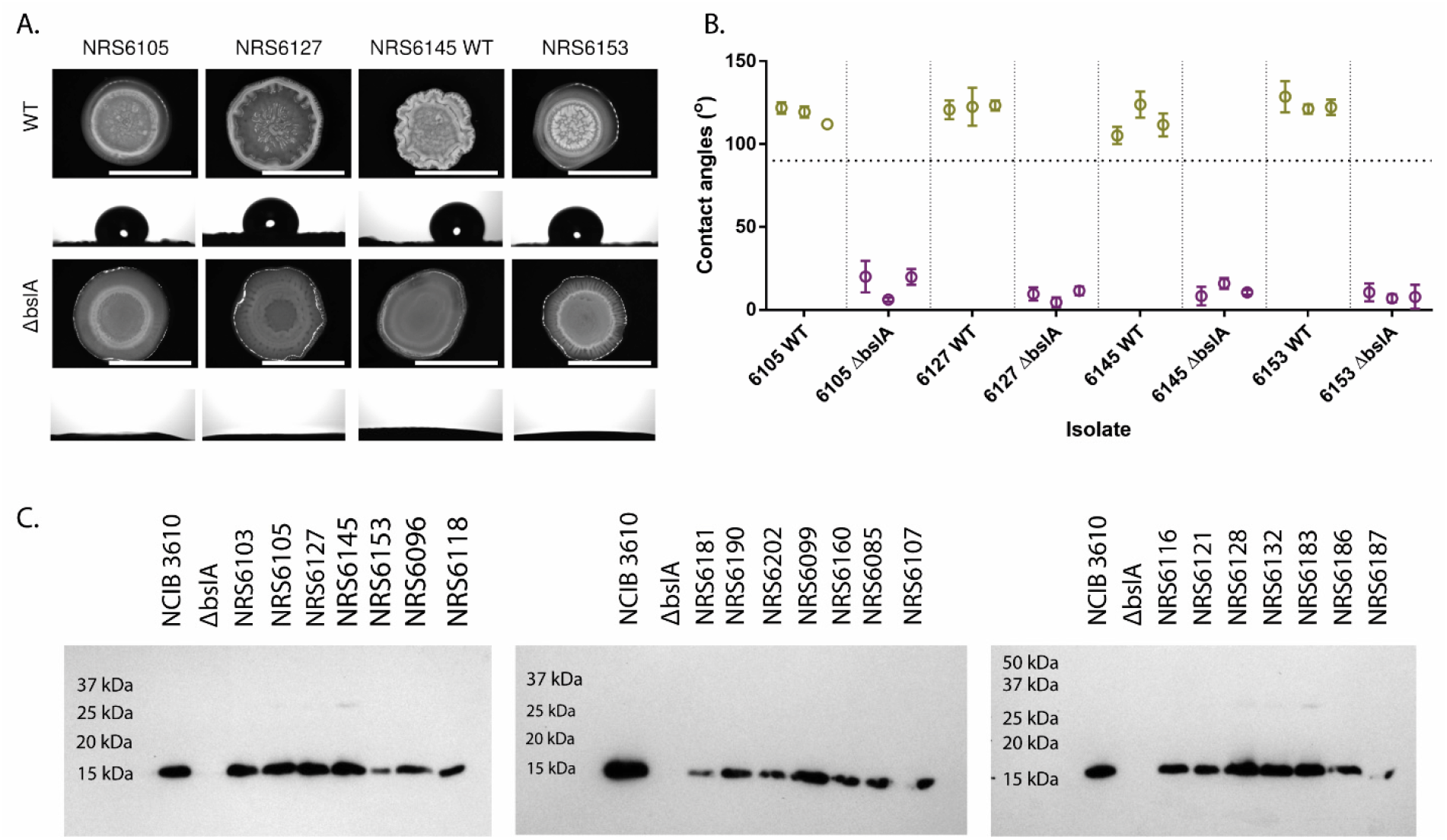
Deletion of *bslA* in hydrophobic environmental isolates of *B. subtilis*. (**A**) Representative images of biofilms formed by the wild type (WT) (top) and *bslA* deletion strains (bottom) of four hydrophobic isolates. Biofilms were grown at 30 °C for 48 h prior to imaging and the scale bars represent 1 cm. The images below the biofilms show a 10 μl droplet of water on the surface of the respective biofilm after 5 min. Scale bars represent 1 cm. (**B**) Biofilm hydrophobicity assay results of wild type (green) and *bslA* mutant variants (purple) of four hydrophobic background strains. The results shown represent three biological repeats per strain, and the three technical repeats are represented at standard deviation error bars on their respective biological repeats. The horizontal line indicates the 90° cut-off value for hydrophobicity, with data points below the line representing a hydrophilic surface and data points above the line indicating biofilm hydrophobicity. The values of the parental strains are the same as shown in Fig. 2C and 4B and are repeated for clarity (Table S5). (**C**) Representative immunoblot analysis of BslA proteins extracted from biofilms grown at 30 °C for 48 h (minimum n=2). The specificity of the antibody is demonstrated by use of the wild type *B. subtilis* isolate NCIB 3610 and corresponding *bslA* mutant (NRS2097). The expected size of monomeric BslA is 14 kDa.

### BslA is produced in biofilms of hydrophobic and hydrophilic isolates

The presence of hydrophobic and hydrophilic isolates in our collection, coupled with the conservation of BslA at the sequence level, led us to hypothesise that BslA may not be produced in the hydrophilic isolates under the conditions used. To test this hypothesis, proteins were extracted from mature biofilm of all the genetically competent isolates, using the reference strain NCIB3610 and the *bslA* negative control strains. Immunoblotting with an anti-BslA antibody revealed the presence of BslA in the mature biofilms from hydrophobic and hydrophilic isolates alike (Figure 3C). Therefore, lack of BslA production is not the reason behind the observed hydrophilicity of some environmental isolates of *B. subtilis*.

### γ-PGA affects biofilm structure of *B. subtilis* isolates

The fact that many of the isolates of *B. subtilis* are not consistently hydrophobic, despite the conservation and production of BslA in mature biofilms, led us to hypothesise that another biofilm matrix exopolymer might be preventing hydrophobicity from manifesting. Poly γ-glutamic acid (γ-PGA) is a hydrophilic polymer that, although it has no impact on biofilm structure in the model isolate NCIB 3610 in the conditions used here (10), has been found to be an important biofilm matrix component in some isolates of the species (10, 11). To test whether γ-PGA could be “masking” hydrophobicity, we constructed deletion mutants of the *pgsB* gene, which encodes part of the biosynthetic machinery that produces γ-PGA (33), in two hydrophilic and two hydrophobic isolates: namely NRS6105 and NRS6153 (hydrophobic) and NRS6069 and NRS6118 (hydrophilic). As expected, the *pgsB* mutants exhibited a dry morphology when grown on LB agar plates (Figure S2). We also found that the structure of the colony biofilm formed by each of the soil isolates was greatly impacted by *pgsB* deletion (Figure 4A). With the strains constructed, we tested hydrophobicity of the colony biofilms. Our hypothesis was that absence of γ-PGA would “reveal” biofilm hydrophobicity in the hydrophilic isolates due to the lack of the water-absorbing polymer. Contrary to our hypothesis, the upper surfaces of both hydrophilic wild type isolates tested remained hydrophilic after deletion of *pgsB* (Figure 4B). Moreover, one of the isolates that formed a hydrophobic upper colony biofilm surface lost biofilm surface hydrophobicity after deletion of *pgsB*. Together these data highlight that absence of γ-PGA has a wider impact on biofilm formation and is likely to interact with other polymeric substances in the biofilm matrix.

**Figure 4:**
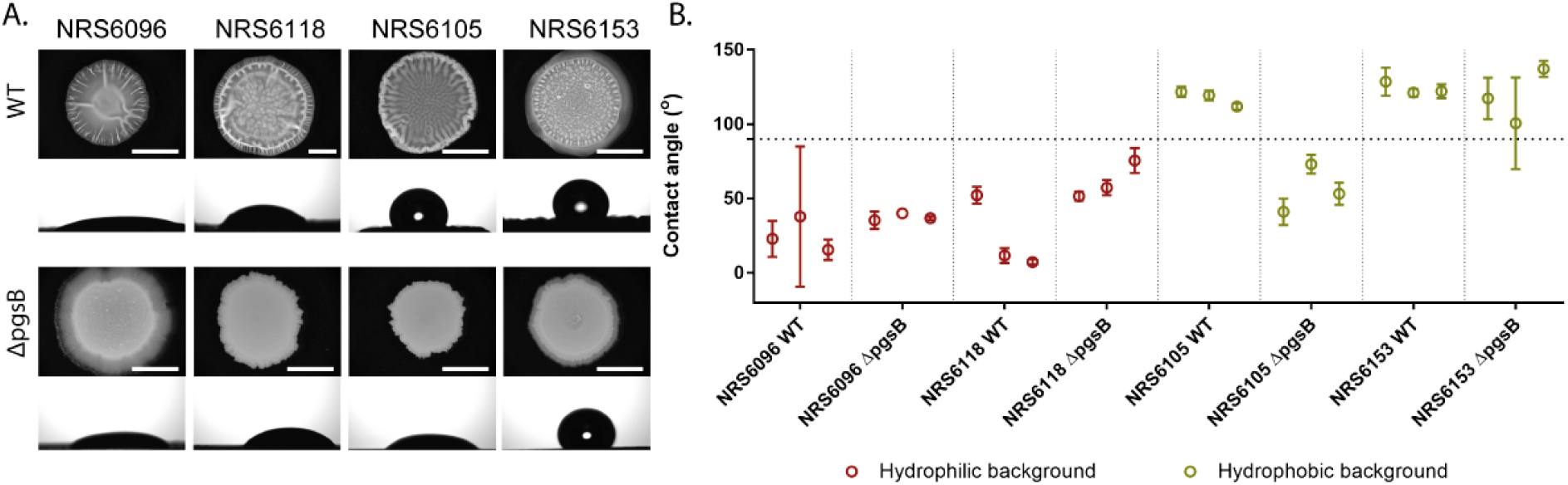
Biofilm morphology and hydrophobicity isolates of WT and *pgsB* mutants of *B. subtilis* soil isolates. (**A**) Representative images of biofilms of wild type (top) and their respective *ΔpgsB* variants (bottom) grown on MSgg media at 30 °C for 48 h. The images below the biofilms show a 10 μl droplet of water on the surface of the respective biofilm after 5 min. Scale bars represent 1 cm. (**B**) Results of hydrophobicity assays of WT and *ΔpgsB* variants of selected environmental isolates of *B. subtilis*. The horizontal line represents the 90° cut-off point for hydrophobicity, such that any points above 90° indicates a hydrophobic surface and points below the 90° line represent a hydrophilic surface. The vertical lines show the separation of the different strains. The three data points correspond to the mean value of each biological replicate. Error bars represent the standard deviation of three technical replicates. Data is coloured by the classification of the parental isolates (WT) as either hydrophilic (green) or hydrophobic (red). The values of the parental strains are the same as show in Fig. 2C and 3B and are repeated for clarity (Table S5).

## Discussion

*Bacillus subtilis* is a diverse species that can colonise many environments and has an open pan-genome (34). Despite this, most research has focused on model isolates, of which NCIB 3610 is predominately used for the analysis of biofilm formation. It is well established that *B. subtilis* NCIB 3610 forms hydrophobic biofilms (8, 15). To test the conservation of biofilm hydrophobicity across a range of *B. subtilis* isolates, we used a citizen science approach to collate a library of environmental isolates. We used these isolates to examine hydrophobicity and found that, of the isolates tested, only 23.8 % (5 of 21) showed a consistently hydrophobic phenotype. Five other isolates formed consistently hydrophilic biofilms and the remaining 11 showed variable or intermediate results. As biofilm hydrophobicity provides a protective mechanism against antimicrobial agents, the variability in overall hydrophobicity appears to be counterintuitive to enhanced survival in biofilms. However, Grau et al., have previously demonstrated through experimental evolution that an isogenic biofilm of the model NCIB 3610 will eventually differentiate into distinct morphotypes when grown in biofilms over multiples generations, some of which are hydrophilic (35). It is therefore possible that the isolates used here, which have been extracted from a natural environment where they are likely to have existed in mixed communities, have diversified into non-hydrophobic variants, despite encoding *bslA*. Consistent with this, the intentional mixing of wild-type isolates of *B. subtilis* that exhibit different surface wetting properties in single culture alters the properties and morphology of the blended isolate community that develops (36).

As both hydrophobic and hydrophilic isolates were present in our collection, we hypothesised that BslA may not be produced in the hydrophilic isolates under the conditions used. However, our results showed that all isolates, hydrophobic and hydrophilic alike, produced BslA, suggesting that the observed biofilm hydrophilicity is not a result of the absence of BslA. It remains to be established if the BslA produced by these isolates is in the form of dimers or monomers (16) since dimerization of BslA is crucial for conferring biofilm hydrophobicity in the model *B. subtilis* NCIB 3610 strain. Dimerization is catalysed by disulphide bond formation between two cysteine residues at the C-terminus (the CxC motif) and is the result of both enzymatic catalysis by thiol-disulphide oxidoreductases and spontaneous oxidation (16). While at a sequence level the CxC motif of BslA is identical in all isolates tested, and therefore dimerization is theoretically possible in all isolates, the localisation of BslA within the matrix could influence the state that the protein is found in. In biofilms, such as those formed by *B. subtilis* on agar surfaces, a steep oxygen gradient forms such that the biofilm surface is an oxygen dense environment, but the oxygen concentration decreases as a function of depth within the biofilm (16). Therefore, a difference in the localisation of BslA to that of NCIB 3610, where the protein migrates to the biofilm-air interface, could result in less dimerization, impacting biofilm hydrophobicity. Correspondingly, if localisation of BslA is impacted and the elastic film of BslA does not form at the air-biofilm interface it would impact biofilm hydrophobicity. Future studies investigating BslA localisation could help uncover the reason behind some isolates having a hydrophilic phenotype despite BslA being produced in mature biofilms.

As mentioned above, while BslA is a “bacterial hydrophobin” (19) and the main protein determinant of biofilm hydrophobicity (8), the presence of other matrix exopolymers (such as the exopolysaccharides) (8) and surface topology (20) impact hydrophobicity of the biofilm. Additionally, there is evidence to suggest that the molecular composition of the matrix varies amongst isolates of *B. subtilis*. γ-PGA has been shown to be the dominant matrix exopolymer in some environmental isolates of *B. subtilis* (10, 11). This contrasts with the model NCIB 3610, where deletion of genomic regions involved in γ-PGA biosynthesis does not have a consistent impact on biofilm architecture (10, 13). γ-PGA is a highly hydrophilic macromolecule, which functions to trap water inside the biofilm and also provides the community with protection from ethanol (37). We therefore questioned if high levels of γ-PGA could mask hydrophobicity mediated by BslA in the hydrophilic isolates. We uncovered that the structure of biofilms was greatly impacted by deletion of *pgsB* in all isolates tested, consistent with reports highlighting the importance of γ-PGA in biofilms of some environmental isolates of *B. subtilis* (10, 11). The absence of γ-PGA did not reveal new hydrophobic properties in the colony biofilms formed by the hydrophilic isolates. In fact, one of the two hydrophobic isolates lost biofilm hydrophobicity after deletion of *pgsB*. Therefore, while these results show that presence of γ-PGA is not responsible for biofilm hydrophilicity, they also reveal the highly variable nature of the biofilm matrix within a species.

## Acknowledgements

Work in the NSW and CEM laboratories is funded by the Biotechnology and Biological Science Research Council (BBSRC) [BB/P001335/1, BB/R012415/1]. M.K. is supported by a Biotechnology and Biological Sciences Research Council studentship [BB/M010996/1]. We are grateful to the Tayport Community Garden, members of the Stanley-Wall lab and the public engagement team at the University of Dundee for their help with the outreach activities. Genome sequencing was provided by MicrobesNG (http://www.microbesng.uk) which is supported by the BBSRC [grant number BB/L024209/1].

## Supplemental Material

**Table S1:** Accession numbers of reference strains.

| Strain | Genbank accession number |
| --- | --- |
| *B. subtilis* subsp. *subtilis* str. NCIB 3610 | GCA_002055965.1 |
| *B. subtilis* subsp. *inaquosorum* str. DE111 | GCA_001534785.1 |
| *B. subtilis* subsp. *spizizenii* str. W23 | GCA_000146565.1 |
| *B. amyloliquefaciens* DSM7 | FN597644.1 |

**Table S2:**
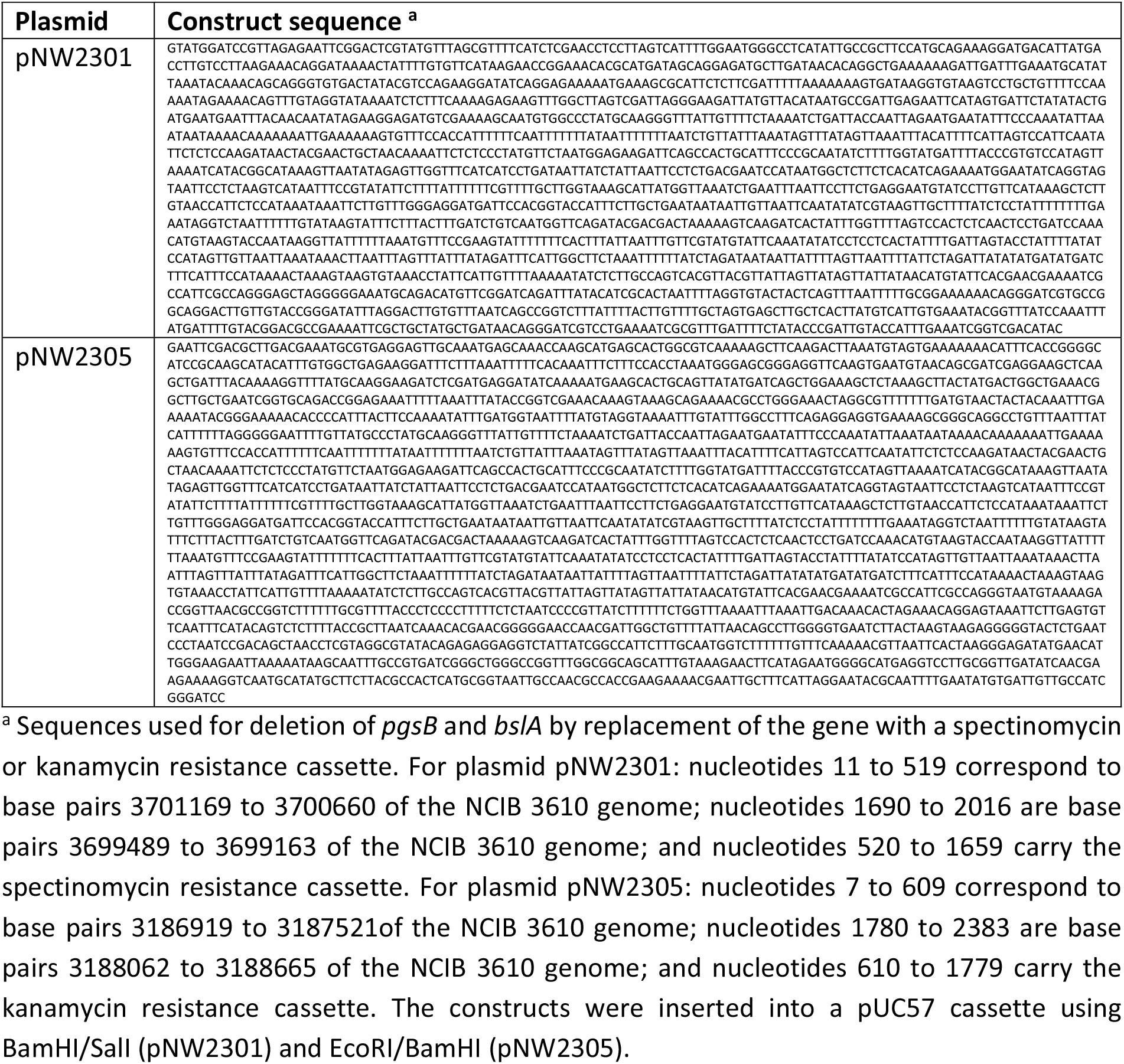
Plasmids synthetically constructed.

**Table S3:** (Preliminary) Classification of isolates of endospore forming bacteria extracted from soil.

| Isolate Number | Preliminary classification <sup>a</sup> |
| --- | --- |
| NRS6072 | *B. mojavensis* |
| NRS6073 | *B. thuringiensis* |
| NRS6074 | *B. megaterium* |
| NRS6075 | *B. cereus* |
| NRS6076 | *B. sp./ B. megaterium* |
| NRS6077 | *B. amyloliquefaciens* |
| NRS6078 | *B. sp. / B. weihenstephanensis* |
| NRS6079 | *B. sp./ B. megaterium* |
| NRS6080 | *B. amyloliquefaciens* |
| NRS6081 | *B. safensis* |
| NRS6082 | *B. sp./ B. altitudinis* |
| NRS6083 | *B. Sp. /B. weihenstephanensis* |
| NRS6084 | *B. subtilis* |
| NRS6085 | *B. subtilis* |
| NRS6086 | *B. sp./ B. cereus* |
| NRS6087 | *Viridibacillus arvi* |
| NRS6088 | *B. sp./ B. cereus* |
| NRS6089 | *B. sp./ B. weihenstephanensis* |
| NRS6090 | *B. sp./ B. megaterium* |
| NRS6091 | *Lysinibacillus fusiformis* |
| NRS6092 | *B. subtilis* |
| NRS6093 | *B. sp./B. thuringiensis* |
| NRS6094 | *B. subtilis* |
| NRS6095 | *B. sp/ B. altitudinis* |
| NRS6096 | *B. subtilis* |
| NRS6097 | *B. sp/ B. safensis* |
| NRS6099 | *B. subtilis* |
| NRS6100 | *B. thuringiensis (looks like B. mycoides)* |
| NRS6101 | *B. flexus* |
| NRS6102 | *B. aryabhattai* |
| NRS6103 | *B. subtilis* |
| NRS6104 | *B. sp./ B. thuringiensis* |
| NRS6105 | *B. subtilis* |
| NRS6106 | *B. thuringiensis* |
| NRS6107 | *B. subtilis* |
| NRS6108 | *B. subtilis* |
| NRS6109 | *B. thuringiensis* |
| NRS6110 | *B. subtilis* |
| NRS6111 | *B. subtilis* |
| NRS6112 | *B. megaterium* |
| NRS6113 | *B. sp./ B. megaterium* |
| NRS6114 | *Viridibacillus arvi* |
| NRS6115 | *B. flexus* |
| NRS6116 | *B. subtilis* |
| NRS6117 | *B. flexus* |
| NRS6118 | *B. subtilis* |
| NRS6119 | *B. megaterium* |
| NRS6120 | *B. subtilis* |
| NRS6121 | *B. subtilis* |
| NRS6122 | *B. megaterium* |
| NRS6123 | *B. megaterium* |
| NRS6124 | *B. sp./ B. megaterium* |
| NRS6125 | *B. megaterium* |
| NRS6126 | *B. sp./ B. megaterium* |
| NRS6127 | *B. subtilis* |
| NRS6128 | *B. subtilis* |
| NRS6129 | *B. flexus* |
| NRS6130 | *B. sp./ B. cereus* |
| NRS6131 | *B. subtilis* |
| NRS6132 | *B. subtilis* |
| NRS6133 | *B. amyloliquefaciens* |
| NRS6134 | *B. subtilis* |
| NRS6135 | *B. sp./ B. weihenstephanensis* |
| NRS6136 | *B. amyloliquefaciens* |
| NRS6137 | *B. subtilis* |
| NRS6138 | *B. flexus* |
| NRS6139 | *B. sp./ B. cereus* |
| NRS6140 | *B. sp. / B. weihenstephanensis* |
| NRS6141 | *B. subtilis* |
| NRS6142 | *Lysinibacillus fusiformis* |
| NRS6143 | *B. megaterium (bad quality)* |
| NRS6144 | *B. flexus* |
| NRS6145 | *B. subtilis* |
| NRS6146 | *B. flexus* |
| NRS6147 | *B. sp./ B. cereus* |
| NRS6148 | *B. subtilis* |
| NRS6149 | *B. pumilus* |
| NRS6150 | *B. sp./ B. altitudinis* |
| NRS6151 | *B. cereus* |
| NRS6152 | *Lysinibacillus fusiformis* |
| NRS6153 | *B. subtilis* |
| NRS6154 | *Lysinibacillus mangiferihumi* |
| NRS6155 | *B. sp. / B. weihenstephanensis* |
| NRS6156 | *B. cereus* |
| NRS6157 | *Lysinibacillus fusiformis* |
| NRS6158 | *B. sp./ B. velezensis* |
| NRS6159 | *B. sp./ B. wiedmannii* |
| NRS6160 | *B. subtilis* |
| NRS6161 | *B. sp./ B. cereus* |
| NRS6162 | *B. sp./ B. amyloliquefaciens* |
| NRS6164 | *B. sp/ B. weihenstephanensis* |
| NRS6166 | *B. flexus* |
| NRS6167 | *B. subtilis* |
| NRS6169 | *Lysinibacillus fusiformis* |
| NRS6170 | *B. flexus* |
| NRS6171 | *B. megaterium* |
| NRS6172 | *Lysinibacillus parviboronicapiens* |
| NRS6175 | *B. sp./ B. cereus* |
| NRS6176 | *B. sp/ B. weihenstephanensis* |
| NRS6177 | *B. cereus* |
| NRS6178 | *B. sp./ B. simplex* |
| NRS6179 | *B. thuringiensis* |
| NRS6180 | *B. sp./ B. altitudinis* |
| NRS6181 | *B. subtilis* |
| NRS6182 | *B. sp./ B. safensis* |
| NRS6183 | *B. subtilis* |
| NRS6184 | *B. flexus* |
| NRS6185 | *B. subtilis* |
| NRS6186 | *B. subtilis* |
| NRS6187 | *B. subtilis* |
| NRS6188 | *B. megaterium* |
| NRS6189 | *B. tequilensis* |
| NRS6190 | *B. subtilis* |
| NRS6191 | *B. sp./ B. safensis* |
| NRS6192 | *B. sp./ B. vallismortis* |
| NRS6193 | *B. sp./ B. megaterium* |
| NRS6194 | *B. subtilis* |
| NRS6195 | *B. sp./ B. amyloliquefaciens* |
| NRS6196 | *B. sp./ B. firmus* |
| NRS6197 | *B. sp./ B. firmus* |
| NRS6198 | *B. amyloliquefaciens* |
| NRS6199 | *B. sp./ B. megaterium* |
| NRS6200 | *B. sp./ B. weihenstephanensis* |
| NRS6201 | *B. sp./ B. simplex* |
| NRS6202 | *B. subtilis* |
| NRS6203 | *B. sp./ B. vallismortis* |
| NRS6204 | *B. subtilis* |
| NRS6205 | *B. subtilis* |
| NRS6206 | *B. subtilis* |
| NRS6207 | *B. methylotrophicus* |
| NRS6208 | *B. sp./ B. amyloliquefaciens* |
| NRS6209 | *B. sp./ B. cereus* |
| NRS6210 | *B. safensis* |
| NRS6211 | *B. licheniformis* |
| NRS6212 | *B. pumilus* |

<sup>a</sup> Species names coloured in black are preliminary classifications based on partial 16S rRNA sequencing whereas those coloured in red are confirmed species classifications based on whole genome sequencing (see Table S4).

**Table S4:** Overview of genome data of the *B. subtilis* isolates.

| Isolate | Contigs | Length (bp) | N50 (bp) | GC (%) | Annotated CDS | Annotated tRNA | Annotated rRNA |
| --- | --- | --- | --- | --- | --- | --- | --- |
| NRS6084 | 49 | 4031155 | 1002034 | 43.62 | 4132 | 84 | 11 |
| NRS6085 | 61 | 4315176 | 678257 | 43.36 | 4397 | 82 | 10 |
| NRS6092 | 59 | 4338566 | 1083827 | 43.27 | 4403 | 86 | 11 |
| NRS6094 | 47 | 4260145 | 665850 | 43.41 | 4322 | 85 | 11 |
| NRS6096 | 50 | 4275968 | 943277 | 43.32 | 4323 | 82 | 10 |
| NRS6099 | 69 | 4074010 | 2123067 | 43.73 | 4060 | 82 | 10 |
| NRS6103 | 60 | 4236010 | 944259 | 43.41 | 4263 | 83 | 11 |
| NRS6105 | 46 | 4061097 | 2041760 | 43.75 | 4012 | 83 | 11 |
| NRS6107 | 67 | 4138021 | 918781 | 43.65 | 4115 | 84 | 11 |
| NRS6108 | 76 | 4420112 | 922980 | 43.28 | 4517 | 84 | 13 |
| NRS6110 | 67 | 4357237 | 608323 | 43.24 | 4417 | 82 | 11 |
| NRS6111 | 51 | 4145435 | 1034199 | 43.53 | 4142 | 83 | 13 |
| NRS6116 | 38 | 4190747 | 918776 | 43.59 | 4196 | 83 | 10 |
| NRS6118 | 62 | 4279571 | 618691 | 43.33 | 4329 | 83 | 11 |
| NRS6120 | 203 | 4038801 | 56365 | 43.2 | 4212 | 84 | 11 |
| NRS6121 | 42 | 4090979 | 2081154 | 43.69 | 4075 | 84 | 11 |
| NRS6127 | 58 | 4198992 | 1049733 | 43.43 | 4239 | 83 | 11 |
| NRS6128 | 37 | 3964538 | 995409 | 43.84 | 3924 | 84 | 11 |
| NRS6131 | 45 | 4210627 | 509484 | 43.16 | 4300 | 82 | 13 |
| NRS6132 | 40 | 4148779 | 944426 | 43.53 | 4153 | 82 | 10 |
| NRS6134 | 41 | 4214740 | 1055486 | 43.5 | 4236 | 85 | 11 |
| NRS6137 | 56 | 4289748 | 851995 | 43.34 | 4359 | 76 | 11 |
| NRS6141 | 48 | 4294788 | 715217 | 43.46 | 4369 | 85 | 11 |
| NRS6145 | 56 | 4148969 | 2098596 | 43.59 | 4112 | 84 | 10 |
| NRS6148 | 61 | 4221073 | 1075194 | 43.43 | 4260 | 84 | 12 |
| NRS6153 | 64 | 4191444 | 2120208 | 43.67 | 4201 | 85 | 11 |
| NRS6160 | 38 | 4188868 | 990131 | 43.49 | 4210 | 82 | 13 |
| NRS6167 | 43 | 4156724 | 531803 | 43.92 | 4106 | 81 | 10 |
| NRS6181 | 65 | 4241152 | 614641 | 43.47 | 4324 | 83 | 10 |
| NRS6183 | 73 | 4225281 | 537639 | 43.41 | 4278 | 80 | 10 |
| NRS6185 | 38 | 4135906 | 1074004 | 43.51 | 4301 | 85 | 12 |
| NRS6186 | 77 | 4299693 | 480191 | 43.28 | 4357 | 82 | 10 |
| NRS6187 | 33 | 4061267 | 1048014 | 43.7 | 4057 | 80 | 10 |
| NRS6190 | 63 | 4081363 | 2087604 | 43.73 | 4050 | 83 | 12 |
| NRS6194 | 128 | 4120078 | 2129494 | 43.85 | 4086 | 83 | 10 |
| NRS6202 | 42 | 4224473 | 918793 | 43.39 | 4264 | 84 | 11 |
| NRS6204 | 45 | 4296050 | 933576 | 43.46 | 4374 | 85 | 11 |
| NRS6205 | 37 | 4293873 | 714950 | 43.45 | 4372 | 85 | 11 |
| NRS6206 | 39 | 4203983 | 974199 | 43.45 | 4253 | 81 | 10 |

**Table S5:** Contact angles of a water droplet placed on biofilms formed by *B. subtilis* isolates.

| Strain | Contact angle ± SD <sup>a</sup> |
| --- | --- |
| NCIB 3610 | 130.85 ± 6.92 |
| NCIB 3610 Δ*bslA* (NRS2097) | 26.07 ± 5.61 |
| NRS6085 | 56.11 ± 30.33 |
| NRS6096 | 24.81 ± 23.39 |
| NRS6099 | 82.27 ± 9.77 |
| NRS6103 | 122.42 ± 4.77 |
| NRS6105 | 115.45 ± 7.01 |
| NRS6107 | 37.16 ± 36.79 |
| NRS6116 | 50.44 ± 50.88 |
| NRS6118 | 23.82 ± 21.83 |
| NRS6121 | 92.43 ± 53.38 |
| NRS6127 | 122.28 ± 6.63 |
| NRS6128 | 59.38 ± 29.91 |
| NRS6132 | 84.09 ± 42.88 |
| NRS6145 | 113.58 ± 10.09 |
| NRS6153 | 123.92 ± 6.92 |
| NRS6160 | 93.74 ± 10.39 |
| NRS6181 | 42.89 ± 23.58 |
| NRS6183 | 95.39 ± 34.16 |
| NRS6186 | 31.32 ± 37.82 |
| NRS6187 | 50.07 ± 32.18 |
| NRS6190 | 8.97 ± 4.25 |
| NRS6202 | 7.72 ± 2.95 |
| NRS6901 | 66.36 ± 23.82 |
| NRS6902 | 121.24 ± 21.34 |
| NRS6903 | 37.47 ± 3.92 |
| NRS6904 | 61.58 ± 11.98 |

<sup>a</sup> The contact angle shown is calculated as the average of three biological and three technical repeats. The error shown is the standard deviation.

**Figure S1:**
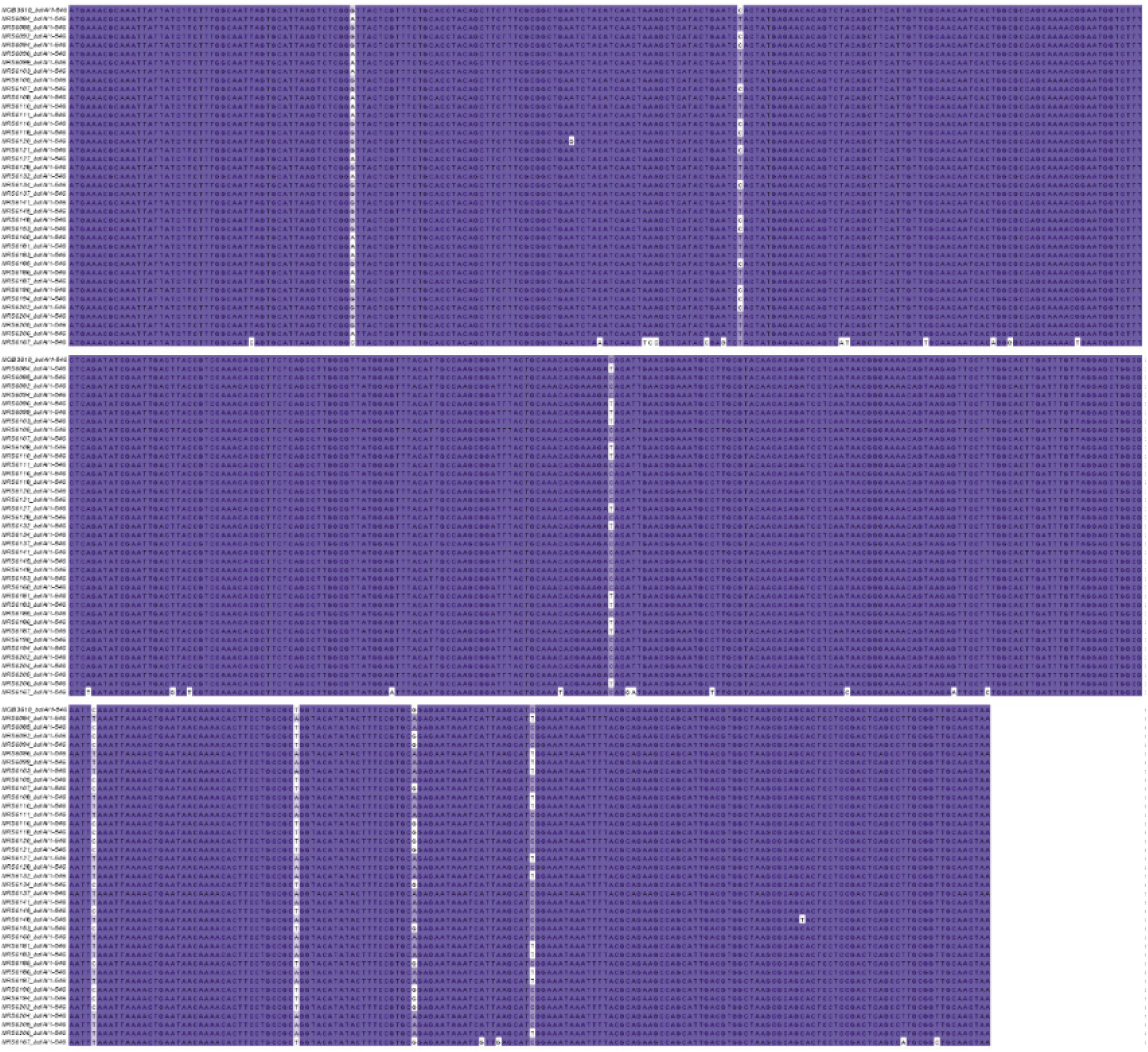
Nucleotide variations in the *bslA* coding region. Alignments of the *bslA* sequences extracted from the whole genome sequencing data of all 39 isolates used in this study. The sequences were aligned in Jalview (1) and the colouring scheme used represents percent identity, where nucleotides coloured in dark blue correspond to an identity of >80% to the consensus sequence. The two lighter shades of blue represent sequence identities of >60% and >40% and nucleotides coloured in white show a sequence identity of less than 40% to that of the consensus sequence.

**Figure S2:**
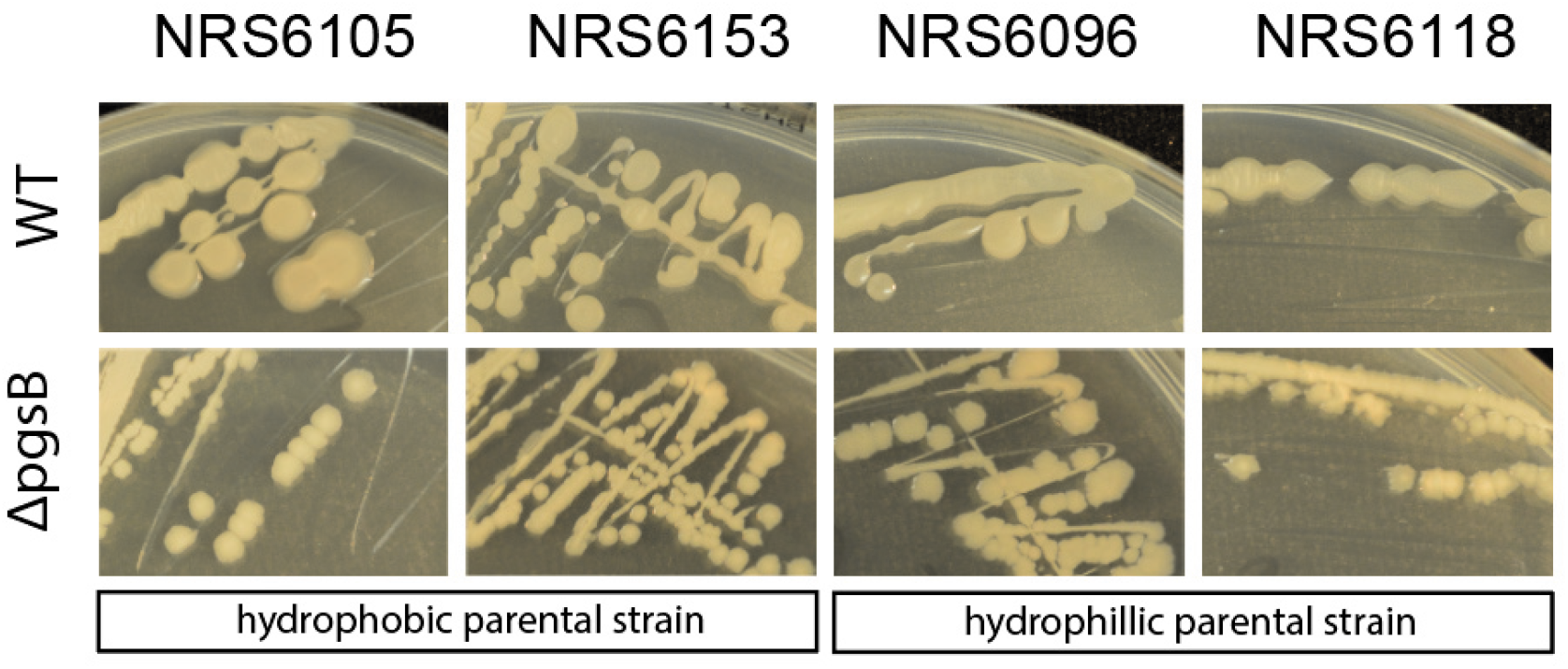
Morphology of WT and *pgsB* mutants of *B. subtilis* soil isolates. Representative images of wild type (top) and their respective Δ*pgsB* variants (bottom) grown on LB media at 37 °C for 16 h.

